# SARS-CoV-2 and its ORF3a, E and M viroporins activate inflammasome in human macrophages and induce of IL-1*α* in pulmonary epithelial and endothelial cells

**DOI:** 10.1101/2023.11.13.566917

**Authors:** Magdalena Ambrożek-Latecka, Piotr Kozlowski, Grażyna Hoser, Magdalena Bandyszewska, Karolina Hanusek, Dominika Nowis, Jakub Gołąb, Małgorzata Grzanka, Agnieszka Piekiełko-Witkowska, Luise Schulz, Franziska Hornung, Stefanie Deinhardt-Emmer, Ewa Kozlowska, Tomasz Skirecki

**Affiliations:** Department of Translational Immunology and Experimental Intensive Care, Centre of Translational Research, Centre of Postgraduate Medical Education, Warsaw, Poland; Department of Molecular Biology, Institute of Biochemistry, Faculty of Biology, University of Warsaw, Warsaw, Poland; Department of Biochemistry and Molecular Biology, Centre of Translational Research, Centre of Postgraduate Medical Education, Warsaw, Poland; Department of Immunology, Medical University of Warsaw, Warsaw, Poland; Institute of Medical Microbiology, Jena University Hospital, Am Klinikum 1, Germany; Department of Immunology, Institute of Functional Biology and Ecology, Faculty of Biology, University of Warsaw, Warsaw, Poland

**Keywords:** caspase-1, respiratory viruses, innate immunity, interleukin-1, pyroptosis

## Abstract

Inflammasome assembly is a potent mechanism responsible for the host protection against pathogens, including viruses. When compromised, it can allow viral replication, while when disrupted, it can perpetuate pathological responses by IL-1 signaling and pyroptotic cell death. SARS-CoV-2 infection was shown to activate inflammasome in the lungs of COVID-19 patients, however, potential mechanisms responsible for this response are not fully elucidated. In this study, we investigated the effects of ORF3a, E and M SARS-CoV-2 viroporins in the inflammasome activation in major populations of alveolar sentinel cells: macrophages, epithelial and endothelial cells. We demonstrated that each viroporin is capable of activation of the inflammasome in macrophages to trigger cell death and IL-1*α* release from epithelial and endothelial cells. Small molecule NLRP3 inflammasome inhibitors reduced IL-1 release but weakly affected the pyroptosis. Importantly, we discovered that while SARS-CoV-2 could not infect the pulmonary microvascular endothelial cells it induced IL-1*α* and IL-33 release. Together, these findings highlight the essential role of macrophages as the major inflammasome-activating cell population in the lungs and point to endothelial cell expressed IL-1*α* as a potential novel component driving the pulmonary immunothromobosis in COVID-19.

## Introduction

Severe acute respiratory syndrome coronavirus 2 (SARS-CoV-2) causing the COVID- 19 disease was responsible for one of the biggest global pandemics that affected millions of people worldwide. The clinical manifestations of COVID-19 ranges from almost asymptomatic disease to acute respiratory distress syndrome (ARDS) characterized by high mortality [1, 2]. Evolution of the virus led to the emergence of subsequent variants that differed in clinical course, infectivity and response to vaccination [3]. Despite tremendous research effort, the exact mechanisms responsible for the control of infection and influencing the severity of the disease remain unclear.

One of the major innate immune mechanisms responsible for early detection and potent response to pathogens, including viruses, is the inflammasome assembly [4]. Activation of the inflammasome sensor proteins leads to formation of macromolecular complex which facilitates autocleavage of pro-caspase-1 into the active form which then cleaves pro-IL-1*β*, pro-IL-18 into active inflammatory cytokines. Moreover, active caspase-1 cleaves gasdermin D into fragments of which the N-terminal one forms pores in cell membrane enabling the release of IL-1 and IL-18 and lead to the lytic cell death termed pyroptosis [4]. In particular, the NLRP3 inflammasome that senses disturbances of cellular homeostasis such as ion efflux, ROS overproduction and lysosome rupture plays a role in the response to multiple viruses including influenza, coronaviruses and SARS-CoV [5]. The involvement of the NLRP3 inflammasome in the pathogenesis of COVID-19 was postulated based on early clinical observations of the increased IL-1 signaling and high lactate dehydrogenase (LDH) levels in severe COVID-19 [1]. Accordingly, several lines of evidence support that hypothesis including presence of the inflammasome activation markers in lungs of deceased COVID-19 patients, including the increased IL-1 and high LDH [6]. However, the role of the inflammasome in the course of the disease remains perplexing; on the one hand, the concentration of inflammasome-related cytokines is higher in more severe cases but on the other hand they do not have predictive value at admission to the intensive care units [7]. Circulating granulocytes from COVID-19 patients have even impaired inflammasome activation capacity [8]. Also, numerous trials with inflammasome inhibitors or anti-IL-1 blockers have failed [7, 9] with the exception of the su-PAR-guided therapy in pre-hospital patients [10]. Ineffective activity of the inflammasome during influenza virus infection can be detrimental [11], while the low activation of the inflammasomes by coronaviruses in bats is suggested to be a tolerance strategy that makes these animals resistant to the development of severe diseases [12].

SARS-CoV activates the human NLRP3 inflammasome via the ORF3a and E viroporins [13, 14]. Also, SARS-CoV-2 is capable of activation the NLRP3 inflammasome in alveolar macrophages [15]. Although it was reported that spike and N proteins are able to activate the inflammasome in human macrophages [16, 17], the effects of viroporins were not thoroughly evaluated. While the involvement of the NLRP3 in the alveolar epithelial cells is disputable, activation of NLRP1 by SARS-CoV-2 was recently demonstrated in this cell type [18]. However, the comprehensive investigation of the potential of SARS-CoV-2 and its viroporins to activate the inflammasome in major lung cell populations was not performed. Therefore, here we aimed to investigate the role of the key SARS-CoV-2 viroporins (ORF3a, E and M) in the major alveolar sentinel cells, including macrophages, epithelial and endothelial cells. Our study shows that macrophages constitute the major inflammasome-activating cell population in the lungs. We also demonstrate that IL-1*α* expressed by endothelial cells is a potential novel component that drives the pulmonary immunothromobosis in COVID-19

## MATERIALS AND METHODS

### Cell culture

THP-1-ASC-GFP cells (InvivoGen, CA, USA) were maintained according to manufacturer’s instructions using RPMI 1640 and 10% FBS, Normocin (50 mg/ml, InvivoGen, CA, USA), 1% Penicillin/Streptomycin (Thermo Fisher Scientific, MA, USA) and Zeocin (100 mg/ml, InvivoGen). U938 WT and NLRP3 KO cells (a kind gift from Thomas Henry [19]), cells were cultured in RPMI 1640 with 10% FBS. HUVEC/TERT2 and HBEC3-KT were purchased from ATCC (VA, USA) and maintained according to manufacturer’s instructions. Human Pulmonary Microvascular Endothelial Cells (HPMEC, Cat. 3000) and Human Pulmonary Alveolar Epithelial Cells (HPAEpiC, Cat. 3200) were obtained from ScienCell Research Laboratories (CA, USA). HEK-293 (ATCC) were cultured in Dulbecco’s Modified Eagle’s Medium (DMEM), 10% FBS heat-inactivated and 1% Penicillin/Streptomycin. Human Small Airway Epithelial Cells (SAECs, Cat. CC-2547) and the corresponding medium were commercially obtained by Lonza (NJ, USA). Human Pulmonary Microvascular Endothelial Cells (HPMECs, Cat. C-12281) and the appropriate medium were obtained from Promocell (Germany). Macrophages were isolated from buffy coats of healthy donors at the Jena University Hospital (ethics approval no. 2019-1519). The blood samples were diluted and separated by using Histopaque 1077 (Sigma-Aldrich, MO, USA) as well as density gradient centrifugation. First, the isolated monocytes were cultured in Monocyte Attachment Medium for 2 hours and then grown in M1-Macrophage Generation Medium DXF for 7 days. All cells were cultured at 37°C in a 5% CO_2_ humidified incubator.

### Transfection of immortalized cell lines

HBEC-3KT cells were transfected with pSELECT-hASC-GFP plasmid (InvivoGen) using linear 25 kDa polyethylenimine (PEI) (Polysciences, PA, USA), with the DNA:PEI ratio was 1:4 (w:w). 2.3x10^5^ cells were seeded on a T25 flask and after 24 hours transfection was performed using 1µg of plasmid. To select stably transfected cells, after 72 hours, selective medium containing Puromycin (4 μg/ml, InvivoGen) was added and after 7 days cells were sorted using FACS Aria (BD, CA, USA). Cells were cultured in selective medium for 3 weeks and medium was changed every 3 days. HUVEC/TERT cells were transfected with pSELECT- hASC-GFP (InvivoGen) plasmid by electroporation as previously described [20] with slight modifications. Suspended cells (1x10^6^) were mixed with 30 μg of plasmid and electroporated at 340 V for 100 μs. For selection of stably transfected cells, medium was replaced with medium containing Zeocin (20 μg/ml, InvivoGen) 96 hours after electroporation. Cells were cultured in selective medium for 3 weeks and medium was changed every 3 days. Then, GFP-positive cells were sorted using FACS Aria (BD, CA, USA).

### Virus strains and infection of the cells

The SARS-CoV-2 wild type, alpha and delta variants were isolated from respiratory specimens of different patients from the Jena University Hospital (ethics approval no. 2018-1263) as previously described [21] (SARS-CoV-2/hu/Germany/Jena-vi005588/2020 (wild type variant), SARS-CoV-2/hu/Germany/Jena-vi043115/2021 (alpha variant) SARS-CoV- 2/human/DEU/vi0114749/2021 (delta variant)). These SARS-CoV-2 variants were cultivated and propagated in Vero-76 cells (ATCC, Cat. CRL-1578) cultured in DMEM supplemented with 10% fetal bovine serum, 1% Penicillin/Streptomycin (Thermo Fisher Scientific), and stocks were titrated via standard plaque assay. All SARS-CoV-2 propagation and infection were conducted in BSL-3 laboratories.

SAECs and HPMECs were either left uninfected (mock) or were infected with SARS-CoV-2 at a multiplicity of infection (MOI) of 1. Therefore, cells were incubated with the appropriate virus-dilution prepared in corresponding medium for 1 hour at 37 °C, 5% CO_2_. After 1 hour, cells were washed with phosphate-buffered saline and were incubated in fresh medium for 1 day or 3 days at 37 °C, 5% CO_2_.

Differentiated macrophages were infected with the SARS-CoV-2 virus variants wild type, alpha and delta. After washing the cells with PBS, they were incubated with the indicated virus diluted in RPMI supplemented with 10 % human serum albumin and 60 minutes later the cells were washed with PBS and cultivated in RPMI containing 10 % HSA and (30 ng/ml) TPCK-treated trypsin. After 8 and 24 hours the supernatants were collected to perform LEGENDplex™

Human Cytokine Panel 2 (BioLegend, San Diego, USA) for cytokine determination and CyQUANT™ LDH assay (Thermo Fisher Scientific) for cytotoxicity detection. For protein analysis, cells were washed with PBS and lysed in ice-cold RIPA buffer.

### Generation of lentiviral constructs encoding SARS-CoV-2 proteins

The plasmids coding for the SARS-CoV-2 E, SARS-CoV-2 M and SARS-CoV2 ORF3a proteins that are tagged with the FLAG epitope on their C-termini (pLV-SARS-CoV-2 E-FLAG, pLV-SARS-CoV-2 M-FLAG and pLV-SARS-CoV-2 ORF3a-FLAG, respectively) were generated using the backbone of pLV-mCherry (a gift from Pantelis Tsoulfas via Addgene, #36084) from which the mCherry sequence was removed by cutting it with the BamHI and SalI restriction enzymes. The coding sequences for the proteins were amplified by PCR with Phusion Hot Start II High-Fidelity DNA polymerase (Thermo Fisher Scientific) using as templates pDONR207 SARS-CoV-2 E, pDONR207 SARS-CoV-2 M and pDONR207 SARS-CoV-2 ORF3a (all plasmids were gifts from Fritz Roth via Addgene, # 141273, #141274 and #141271, respectively) [22] and the pairs of primers, 5’-GCGGGATCCGCCGCCACCATGTACTCTTTCGTGAGC-3’ and 5’-CGTCATCCTTGTAATCGCCCACCAGCAGGTCAGGCAC-3’, 5’-GCGGGATCCGCCGCCACCATGGCTGACTCTAACGGT-3’ and 5’-CGTCATCCTTGTAATCGCCCTGCACCAGCAGGGCGAT-3’, and 5’-GCGGGATCCGCCGCCACCATGGACCTGTTCATGAGA-3’ and 5’-CGTCATCCTTGTAATCGCCCAGTGGCACGGAGGTGGT-3’, respectively (the BamHI sites are underlined). To obtain the full-length FLAG-tagged constructs, each PCR product was reamplified with the specific forward primer (with the BamHI site) and the common reverse primer, 5’-CGC<u>GTCGA</u>CTTACTTATCGTCGTCATCCTTGTAATCGC-3’ (the SalI site is underlined). The final PCR products were cut with BamHI and SalI, ligated with the pLV backbone using T4 DNA ligase and the NEB Stable Competent Escherichia coli host cells (#C3040H, New England Biolabs, MA, USA) were transformed. Plasmid DNAs were isolated and all sequences coding for the proteins were positively verified by DNA sequencing.

### Lentivirus generation

For lentivirus production, 1.5 × 10^6^ HEK-293T cells were seeded into 10 cm dishes 72 h prior to transfection in 10 ml of DMEM supplemented with 10% FBS (Invitrogen) and P/S solution (Thermofisher). The lentiviruses encoding the gene of interest (GOI) were introduced to the packaging HEK293T cells by calcium phosphate co-transfection with 8.6 μg of a GOI- containing vectors (pLV-SARS-CoV-2-E-FLAG, pLV-SARS-CoV-2-M-FLAG and pLV-SARS-CoV-2-ORF3a-FLAG) and components of 2^nd^ generation of packaging vectors: 8.6 μg of psPAX2 packaging vector and 5.5 μg of pMD2.G envelope vector. Right before transfection HEK293T culture medium was exchanged to 5 ml of fresh DMEM + 10% FBS w/o P/S. The plasmids were resuspended in 450 μl of 300 mM CaCl_2_ and added dropwise to 450 μl of 2 × concentrated HEPES-buffered saline (280 mM NaCl, 20 mM HEPES, 1.5 mM Na_2_HPO_4_, 10 mM KCl and 12 mM D-glucose pH7.2) by vortexing. The precipitate was immediately added to the cell culture medium with gentle swirling. The medium was replaced with 6 ml of fresh DMEM + 10% FBS + P/S 16 hours post transduction. Virus-containing supernatants were collected 48 hours later, centrifuged for 5 minutes at 300 × g, filtered through 0.45 μm low-protein-binding filters (Millipore) and concentrated via 16 hours centrifugation at 3000 x g at 10 °C. The titer of viral particles was estimated using a Lenti-X p24 Rapid Titer Kit (Clontech) according to the manufacturer’s protocol. MOI was calculated by dividing the vector titer for the number of cells transduced.

### Lentiviral infection

The day before infection THP-1-ASC-GFP cells were seeded at 4 x 10^4^ cells/well in a 96- well plate and differentiated with Phorbol 12-myristate 13-acetate (20 ng/ml). After 4 h of PMA treatment medium was replaced for Opti-Mem Reduced Serum Medium without phenol with 5% FBS. On infection day half of the medium was removed, and 30 minutes prior to transduction cells were treated with NATE and different inhibitors were added including: MCC950 (10 μM; Bio-techne Brand), Colchicine (10 μM; Sigma) and Disulfiram (20 μM; Sigma), KCL (50 mM; Sigma). All lentiviral particles: pLV – M, pLV – E, pLV – ORF3a and pLV-mCherry were added at MOI 3 in Opti-Mem medium supplemented with Polybrene (8 μg/ml). Following infection cells were spinoculated at 2000 x g in 32 °C for 1 hour. After 24 hours of infection, medium was collected and stored at -80°C for further cytokine assays.

For infection of HBEC-3KT-ASC-GFP and HPAEpiC the cells were seeded at 8 x 10^3^ cells/well the day before lentiviral infections. HUVEC-ASC-GFP and HMPEC cells were seeded at 1 x 10^4^ cells/well in 96-well plates. On infection day half of the medium was removed, and 30 minutes prior to infection cells were treated with different inhibitors. All lentiviral particles were added to the cells at MOI 3 in corresponding medium supplemented with Polybrene (4 μg/ml).

### Transfection of HEK-293

The transfection was performed using jetOptimus transfection reagent (Polyplus, France, Cat. 101000051). Briefly on day 0 HEK-293 cells were seeded on coverslips covered with poly-L-lysine hydrobromide at 8 x 10^4^ cells/well in a 24-well plate. On day 1 the growth medium was replaced with fresh one without antibiotics. 0.5 µg of each plasmids: pLV-M, pLV-E, pLV- ORF3a and pLV-mCherry were mixed in 50 µl of jetOptimus buffer, vortex and jetOptimus reagent was added at ratio 1:2. Following 10 minutes the transfection mixtures were added to the cells. After 4 hours of incubation the medium was changed to a growth medium. The next day half of the medium was replaced. 72 hours after transfection immunocytochemistry was performed. For immunoblot analysis HEK-293 were seeded at 1.4 x 10^4^ cells/well in a 12-well plate. Next day cells were transfected with mixture: 1 µg DNA, 100 µl jetOptimus buffer, and jetOptimus reagent at 1:2 ratio. On the third day after transfection the cells were washed with cold PBS and lysed in a cold RIPA Lysis Buffer supplemented with Protease Inhibitor Cocktail and 1 mM PMSF. Cell lysates were collected and frozen at -80 °C.

### Western blot

Lysates from transfected HEK-293, infected: macrophages, epithelial and endothelial cells were mixed in 4x Laemmli Sample Buffer supplemented with b-mercaptoethanol and boiled for 5 minutes at 94 °C or heated for 40 minutes at 37 °C for GSDMD immunodetection. 10 µg of proteins were separated in 12.5 % SDS – PAGE gels and blotted onto nitrocellulose or PVDF membranes by semi-wet transfer for 10 minutes at 2.5 A and 25 V. Membranes were blocked in TBS-T with 5 % non – fat milk for 1 hour in RT following incubation overnight at 4 °C with primary (1:1000) antibodies: Flag (Sigma, Cat. F1804), IL-1β (Cell Signaling Technology, MA, USA, Cat. 2022 and Affinity Biosciences, Ohio, USA, Cat. AF4012), IL-1α (Santa Cruz, CA, USA, Cat. sc-9983), NLRP3 (Adipogen, Switzerland, Cat. AG-20B-0014-C100), GSDMD (Cell Signaling Technology, Cat. 96458), β-actin (Novus, CO, USA, Cat.NB600), GAPDH (Millipore, MA, USA, Cat. MAB374; 1:5000). Appropriate mouse or rabbit HRP-conjugated secondary antibody (Vector Laboratories Inc., CA, USA, Cat. PI-2000, PI-1000; 1:10000) was used to detect primary antibodies. Signals were developed by *WesternBright^®^ ECL* HRP substrate (Advansta, CA, USA) and visualized using UVITEC gel documentation system. Proteins band densitometry was done using Image Lab Software (Bio-Rad, CA, USA).

### Immunocytochemistry

Experiments for immunofluorescence staining were performed in 24-well plates and SAECs and HEK-293 cells were cultured on coverslips. After infection, cells were fixed with 4 % paraformaldehyde (PFA) in PBS for at least 30 min at 37 °C and permeabilized with PBS containing 0.1 % Triton-X for 5 minutes and blocked with blocking solution (3 % BSA in PBS) at RT for 30 minutes (all reagents from Sigma-Aldrich). Then, coverslips were incubated with different primary antibodies: anti-SARS-CoV-2 spike antibody (GeneTex, CA, USA, Cat. GTX635807; 1:500), anti-dsRNA monoclonal antibody (Jena Bioscience, Germany, Cat. RNT-SCI-10010200; 1:500) and anti-Flag (Sigma, Cat. F1804; 1:200) diluted in blocking solution at 4 °C overnight. Afterwards, the appropriate antibodies: Alexa Fluor® 488-conjugated donkey anti-rabbit (Jackson ImmunoResearch, PA, USA; 1:500), Cy-3 conjugated donkey anti-mouse (Jackson ImmunoResearch; 1:500) and Alexa Fluor™ 555-conjugated donkey anti-mouse (Invitrogen; 1:500) were incubated for 1 hour at room temperature. Alexa Fluor™ 647 Phalloidin (Thermo Fisher Scientific, Cat. A30107; 1:500) was applied together with secondary antibodies for the visualization of cell borders. Finally, cover slips with stained cells were mounted with DAPI Fluoromount-G (Southern Biotech, AL, USA, Cat. 0100-20) on microscope slides and analyzed by an AxioObserver Z.1 Microscope with 40x or 63x objectives (Zeiss, Jena, Germany).

### Cytokine assays

LDH-Glo Cytotoxicity Assay (Promega, WI, USA, Cat. J2380) was performed according to manufacturer instructions. Briefly the supernatants from treated sample wells were collected and 50x diluted in LDH Storage Buffer and frozen at –20 °C. Then 12.5 μl of thawed samples were incubated with equal volume of LDH Detection Reagent for 1 hour at room temperature before recording luminescence level. To calculate cytotoxicity of lentivirus proteins on THP- 1-ASC-GFP, endothelial and epithelial cells Maximum LDH Release Control for high background was created by adding 10 % Triton X-100 to the cells. Culture medium without cells served as Minimum LDH Release Control. Standard curve was performed for LDH positive control to establish that RLU values of experimental samples are within the linear range of the assay. The assay was performed in white half area 96-well plates (PerkinElmer) in Synergy H2 microplate reader. Lumit Human IL-1β Immunoassay (Promega) was performed according to manufacturer instructions. Briefly 12.5 μl of serially diluted standard and culture medium from infected THP1-ASC-GFP, endothelial and epithelial cells were transferred to the assay plate and mixed with 12.5 μl of 2x antibody mixture. Following 60 minutes incubation at 37 °C 12.5 μl of Lumit Detection Reagent B was added to the samples and 1L-1β release was measured. The assay was performed in 96 – half area white plates in Synergy H2 (Biotek, USA) microplate reader.

### Time-lapse fluorescence microscopy

For performing live-cell fluorescence imaging of infected macrophages, epithelial and endothelial cells IncuCyte® SX1 Live-Cell Analysis System (Sartorius, Germany) was used with green (ex. 440-481 nm/em. 503-544 nm) and red channel (ex.567-607 nm/em. 622-704 nm). Fluorescent signals were monitored in real-time with a microplate reader for 24 h within a tissue culture incubator (37 °C, 5 % CO_2_) on black OptiPlate 96-well (PerkinElmer, MA, USA) under 20x objective. Cells cytotoxicity (Red object counts) were assessed using DRAQ7 (Invitrogen, Cat. D15105) staining, ASC-specs formation (Green object count) was calculated using IncuCyte SX1 analysis system.

### Statistical analysis

Statistical analysis was performed with GraphPad Prism 6 software (GraphPad, CA, USA). All experiments were performed at least in triplicate. Differences between two groups were analyzed using a Student’s *t* test. Multiple group comparisons were performed using two-way ANOVA followed by Bonferroni post hoc test or Kruskal-Wallis test for non-parametric data. The bar graphs represent mean ± standard error. P-values less than 0.05 were considered statistically significant.

## RESULTS

### SARS-CoV-2 ORF3a, E and M viroporins activate NLRP3 inflammasome in human macrophage cell-lines

To analyze the effects of the specific viroporins on inflammasome activation in macrophages, we selected ORF3a, E and M proteins that were previously reported in SARS-CoV to act as ion channels or disrupt intracellular membranes that could lead to inflammasome activation [13, 14]. Firstly, using lentiviral expression system, we transduced human macrophages with genes encoding these proteins (**Fig. 1 A**) and confirmed that they reach different subcellular localizations (**Fig. 1 B**). Then, we took the advantage of the time-lapse fluorescent microscopy to evaluate the effects of SARS-CoV-2 viroporins on the kinetics of cell death and inflammasome assembly in the reporter THP-1-ASC-GFP cell line. Indeed, production of each of the tested viral proteins increased NF-*κ*B-dependent expression of the ASC::GFP fusion protein and formation of the ASC specks indicative of the inflammasome formation (**Fig. 2 A**). Concomitantly, cell death of THP-1 cells was observed measured by the increase in membrane impermeable DRAQ7 nuclear dye uptake in cells infected with lentiviruses carrying the plasmids with viroporins, but not cells infected with control lentiviruses (**Fig. 2 A**). Both events, formation of the ASC specks and cell death were induced quickly after expression of viral proteins. ORF3a protein showed the most potent activity leading to death and disintegration of most of the cells within 10 hours (**Fig. 2 A, B**). The ORF3a and E proteins, but not M protein induced secretion of IL-1*β* which is another hallmark of the inflammasome activity (**Fig. 2 C**).

**Figure 1.**
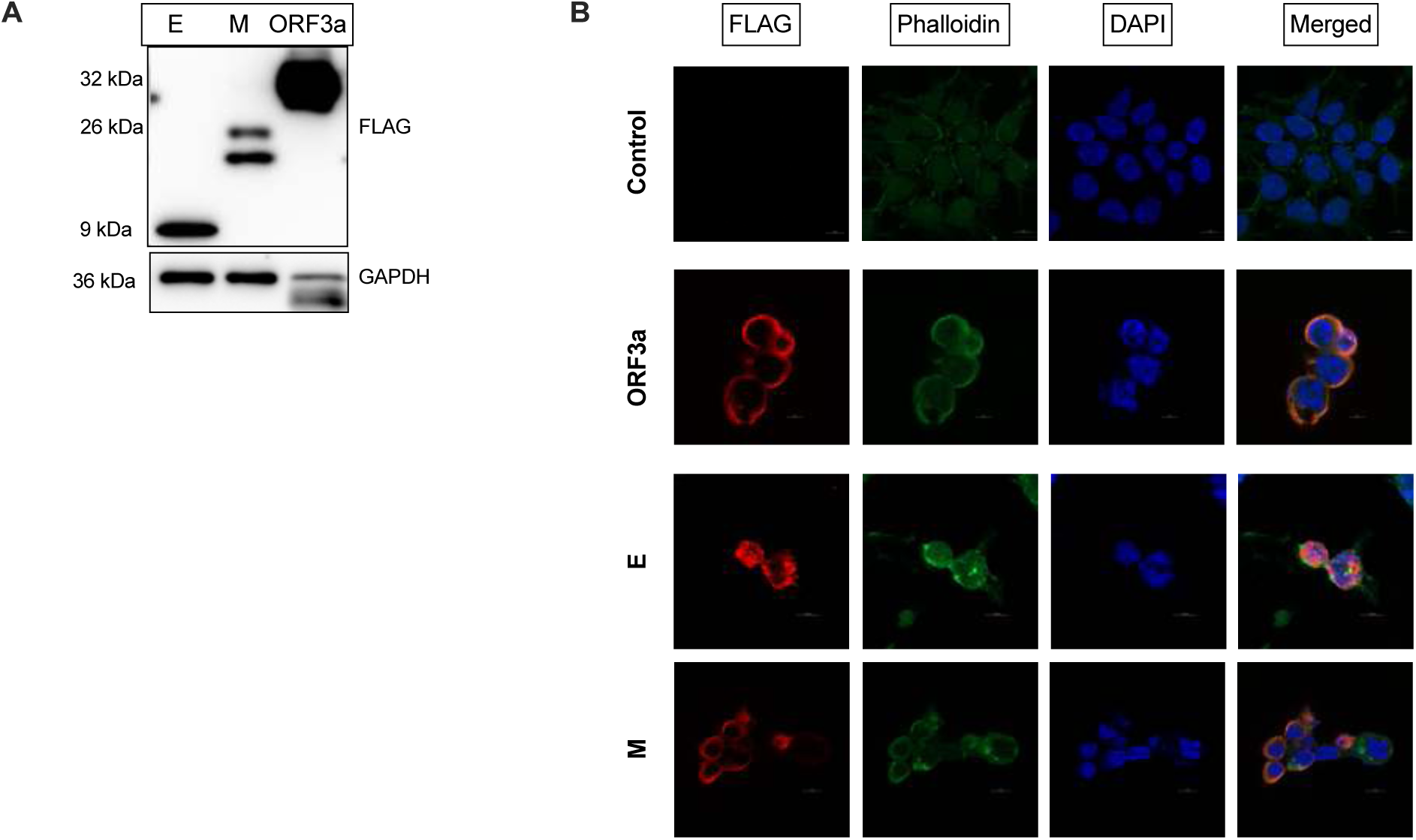
Human cells express the SARS-CoV-2 proteins: ORF3a, E and M upon infection with lentiviruses containing given constructs. A. Western Blot analysis of the FLAG-tagged viral proteins expressed in HEK-293 cells confirm their molecular weight. B. Immunofluorescence of anti-FLAG staining of HEK-293 cells show different subcellular localization of the SARS-CoV-2 viroporins.

**Figure 2.**
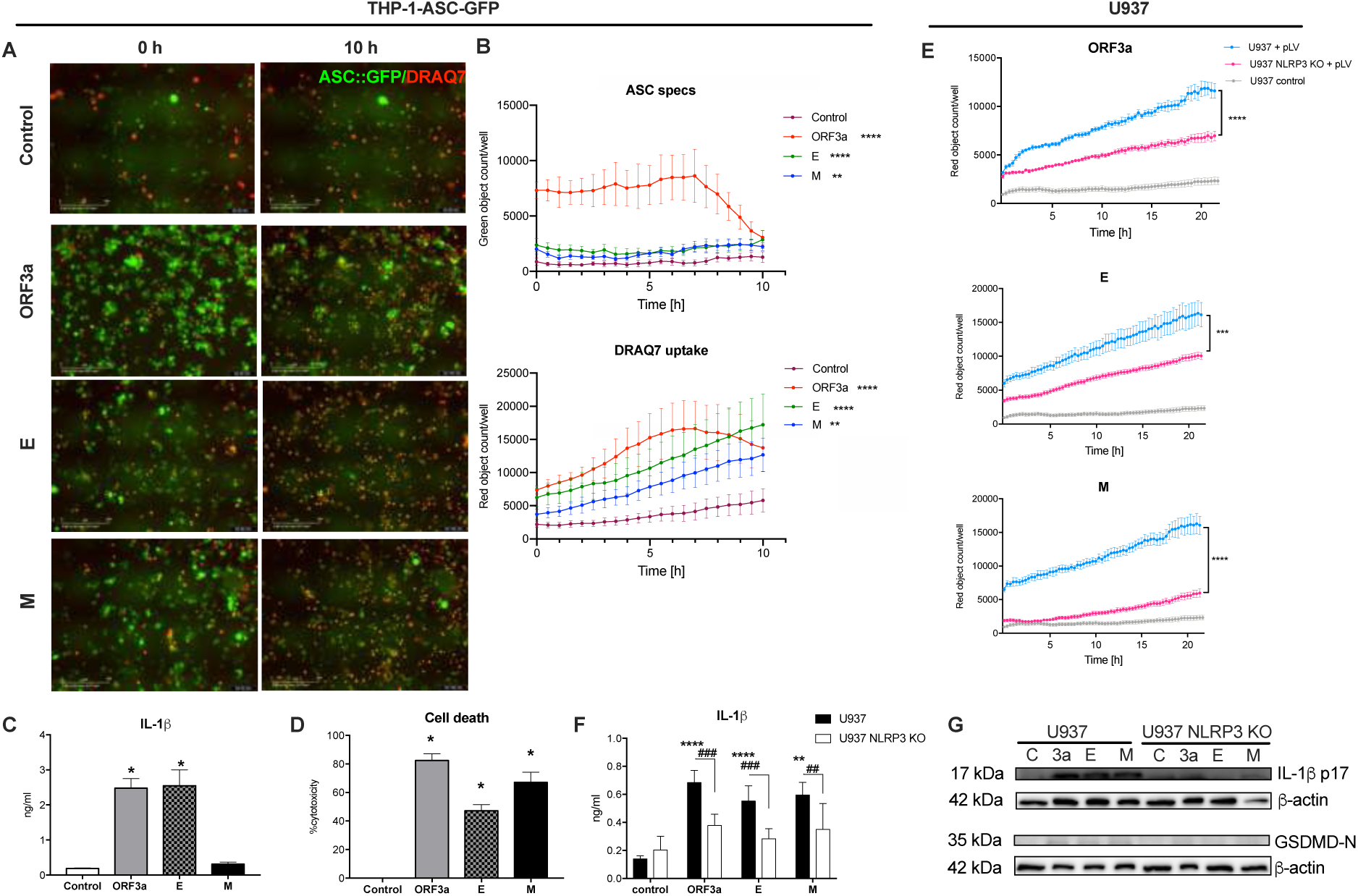
SARS-CoV-2 viroporins activate inflammasome in human macrophages. THP- 1-ASC-GFP reporter cells were infected with lentiviral particles with expression constructs for the viroporins and then observed for the assessment of inflammasome activation. Time-lapse analysis revealed induction of cell death (red, DRAQ7) and ASC-containing inflammasome assembly (green specks) by all tested viroporins. A. Representative pictures (x20 magnification). B. Comparison of kinetics of the ASC speck formation and number of dead cells. C. Analysis of IL-1*β* secretion 24 hours post-infection. D. Cytotoxicity was also measured 24 hours post-infection by the LDH release. E. Analysis of cell death of U937 WT and NLRP3 KO cell lines in response to expression of the viroporins. F. IL-1*β* release from U937 and U937 NLRP3 KO cell lines upon viroporins expressions. G. Immunoblotting of the lysates of U937 cells shows requirement of NLRP3 for cleavage of IL-1*β* and GSDMD. Graphs represent means ± SEM, n=3-5, multiple comparisons were performed with Kruskall-Wallis test. p<0.05 vs control is marked: *p<0.05, **p<0.01, ***p<0.001, ****p<0.0001.

In line with cell death was evident by fluorescent microscopy, we confirmed high level of cytotoxicity in viroporin-treated macrophages by the mean of the release of LDH (**Fig. 2 D**). SARS-CoV-2 viroporins can potentially activate the NLRP3 inflammasome by triggering disturbances in the cellular homeostasis. Therefore, we compared pyroptosis induction by the tested proteins in wild-type (WT) human U937-derived macrophages and NLRP3 KO cells. All three viroporins induced massive pyroptosis in WT cells, while NLRP3 deficiency significantly reduced this effect, yet did not block it completely (**Fig. 2 E**). NLRP3 was also required for the release and maturation of the IL-1*β* (**Fig. 2 F, G**) and GSDMD cleavage upon viroporin expression (**Fig. 2 G**). Altogether, these findings indicate that SARS-CoV-2 ORF3a, E and M proteins activate inflammasome in human macrophages, which at least in part depends on the NLRP3 pathway.

### NLRP3 inflammasome inhibitors reduce IL-1*β* secretion but not cell death upon viroporins expression in human macrophages

Inhibition of the NLRP3 inflammasome was suggested as one of the therapeutic strategies in both pre-clinical and clinical trials [23]. For this reason, we tested under controlled conditions the efficacy of the selected NLRP3 inhibitors to block the viroporin-induced inflammasome activation in human macrophages. The specific NLRP3 inhibitor MCC950 unexpectedly increased ASC speck formation by ORF3a and cytotoxicity by protein E (**Fig. 3 A, B**). Colchicine, which inhibits the NLRP3 inflammasome assembly [24] did not reduce speck numbers upon expression of all three viroporins, as it did not inhibit pyroptosis which was even augmented by this treatment (**Fig. 3 A, B**). In line with these observations, colchicine treatment did not prevent increased LDH release induced by ORF3a and M proteins. In contrast, colchicine attenuated LDH release induced by protein E (**Fig. 3 C**). Disulfiram which is a gasdermin D inhibitor decreased ASC speck formation by ORF3a and E proteins (**Fig. 3 A),** however it did not reduced cytotoxicity significantly (**Fig. 3 B**). Finally, KCl which by prevention of the potassium efflux can inhibit the NLRP3 inflammasome activation neithert reduced speck formation upon ORF3a and M treatment (**Fig. 3 A**) nor attenuated cytotoxicity (**Fig. 3 B, C**). In contrast, all tested inhibitors diminished release of IL-1*β* induced by ORF3a and E proteins, while MCC950 and colchicine reduced M protein - induced release of IL-1*β* (**Fig. 3 D**). These findings suggest that in macrophages cell death in response to the SARS-CoV-2 viroporins is uncoupled from the IL-1*β* release and none of the tested small molecule inhibitors was able to prevent these effects in response to each of the viroporins.

**Figure 3.**
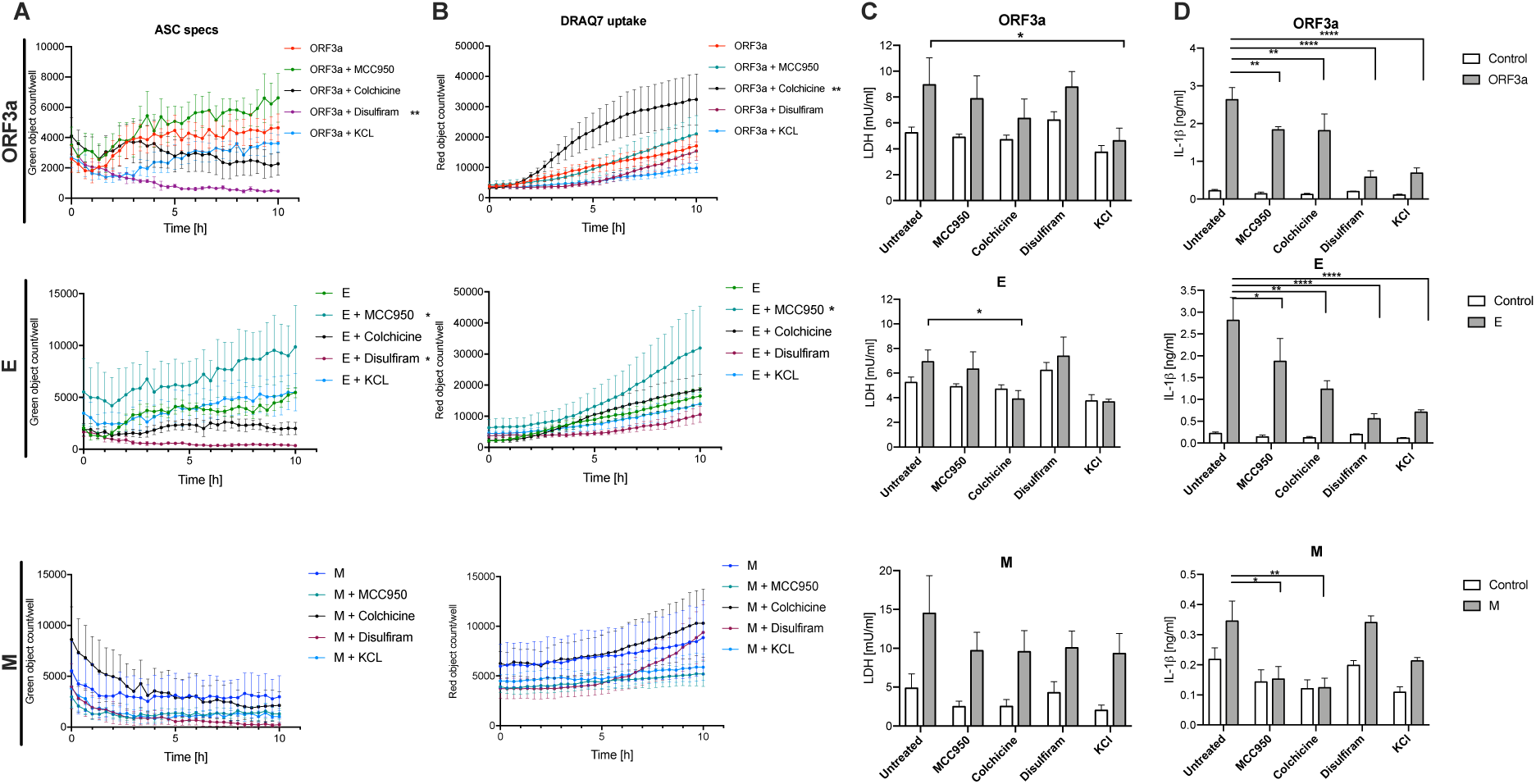
Analysis of the efficacy of the selected NLRP3 inflammasome inhibitors in human THP-1 macrophages expressing SARS-CoV-2 viroporins. SARS-CoV-2 proteins ORF3a, E and M were expressed in THP-1-ASC-GFP cell line to trace inflammasome assembly and pyroptosis in the presence of selected inhibitors of the inflammasome: MCC950, colchicine, disulfiram and KCl. A. Time-lapse analysis of ASC-speck formation. B. Time-lapse analysis of cell death kinetics. C. Level of cell death assessed by LDH secretion 24 hours after lentiviral infection. D. IL-1*β* secretion 24 hours after lentiviral infection. Graphs represent means ± SEM, n=3-5, multiple comparisons were performed with Kruskall-Wallis test. p<0.05 vs control is marked: *p<0.05, **p<0.01, ***p<0.001, ****p<0.0001.

### SARS-CoV-2 viroporins induce cell death in immortalized human bronchial epithelial cells and human umbilical cord endothelial cells without formation of the ASC-dependent inflammasome

As both epithelial and endothelial cells are the major cell type populations in the lungs and both can be infected with SARS-CoV-2 we aimed to investigate whether the viroporins can activate NLRP3 inflammasome in these cell types. In order to be able to perform real-time kinetic assay we first generated reporter cell lines expressing the ASC::GFP fusion protein using bronchial epithelial (HBEK-3KT) and endothelial (HUVEC/TERT2) cell lines. Both generated reporter cell lines, HBEK-3KT-ASC-GFP and HUVEC/TERT2-ASC-GFP were responsive to model NLRP3 inflammasome activation by LPS and nigericin (**Suppl. Fig. 1**). Expression of neither of the SARS-CoV-2 viroporins induced formation of ASC specks in the epithelial nor endothelial cells (**Fig. 4 A, D,** respectively). However, ORF3a, E and M viroporins triggered cell death in both cell types (**Fig. 4 A-F**). Intriguingly, M protein had the weakest effect in epithelial cells, while it evoked the most potent cytotoxicity in endothelial cells. MCC950 was able to reduce ORF3a- and E-induced cell death in epithelial cells. In endothelial cells, only cell death induced by protein M was reduced by MCC950 (**Fig. 4 B, E**). These results indicate that the SARS-CoV-2 viroporins can induce cell death in the epithelial bronchial and endothelial cells that is only partially dependent on the NLRP3 inflammasome and most likely the mode of cell death differs between these cell lines.

**Figure 4.**
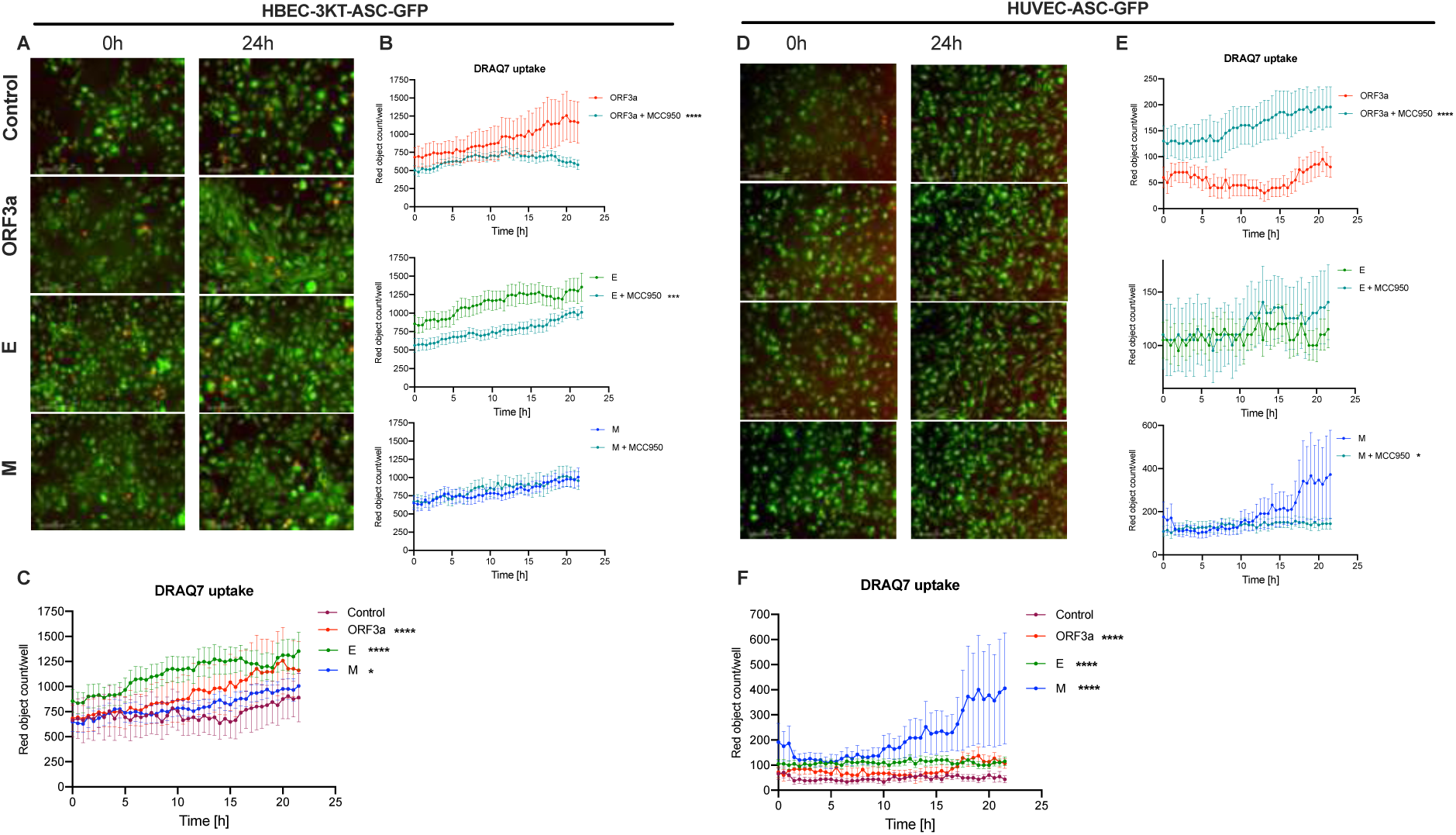
Real-time evaluation of the inflammasome activation in human bronchial epithelial and umbilical vein endothelial cells upon SARS-CoV-2 viroporin expression. Immortalized human bronchial epithelial cells and human umbilical vein endothelial cells expressing the ASC::GFP fusion protein (HBEK-3KT-ASC-GFP, HUVEC-ASC-GFP, respectively) were infected with lentiviral particles to express the SARS-CoV-2 proteins ORF3a, E and M. Green fluorescence represents ASC::GFP and red represents DRAQ7- incorporating dead cells. Representative pictures, comparison of kinetics of cell death upon expression of different viroporins and the effects of MCC950 on cell death are shown for: A., B., C. HBEC-3KT-ASC-GFP. C., D., E. HUVEC-ASC-GFP. Graphs represent means ± SEM, n=3. Multiple comparisons were performed with two-way ANOVA or Kruskall-Wallis test. *p<0.05, **p<0.01, ***p<0.001, ****p<0.0001.

### Neither primary human alveolar epithelial nor endothelial cells activate inflammasome in response to SARS-CoV-2 viroporins

As the reporter cell lines are undifferentiated and immortalized cells, which may differ in various aspects from the primary cells, we decided to evaluate the capacity of SARS-CoV-2 viroporins to activate inflammasome in human pulmonary alveolar epithelial cells (HPAEpiC) and human pulmonary microvascular endothelial cells (HPMEC). Of the proteins tested, the ORF3a and M expression induced release of LDH from HPAEpiC that was accompanied by the increase in cell membrane permeability to DRAQ7 (**Fig. 5 A**). E protein increased membrane permeability but did not induce a significant release of LDH (**Fig. 5 A**). Also, ORF3a and E viroporins induced HMGB1 release (**Fig. 5 B**), but the activity of caspase-1 was not increased by expression of neither of the viroproteins (data not shown). Therefore, we analyzed activation of caspase-3/7 which mediate apoptotic cell death pathway using fluorescent probe. No signal of the pro-apoptotic caspases’ activation was observed in HPAEpiC (**Suppl. Fig. 2 A**). In HPMEC, ORF3a expression also triggered LDH release and DRAQ7 incorporation (**Fig. 5 C**) followed by increase in the HMGB-1 release (**Fig. 5 D**). Furthermore, analysis of caspase- 3/7 revealed that ORF3a triggered their activation in HPMEC (**Suppl. Fig 2 B**). In addition, we analyzed the expression and maturation of inflammasome-related proteins in HPMEC. Expression of ORF3a and M viroporins increased maturation of IL-1*α* and the latter induced also IL-1*β* (**Fig. 5 E**). These experiments revealed capacity of primary pulmonary microvascular endothelial cells to undergo cell death and IL-1 release by SARS-CoV-2 viroporins.

**Figure 5.**
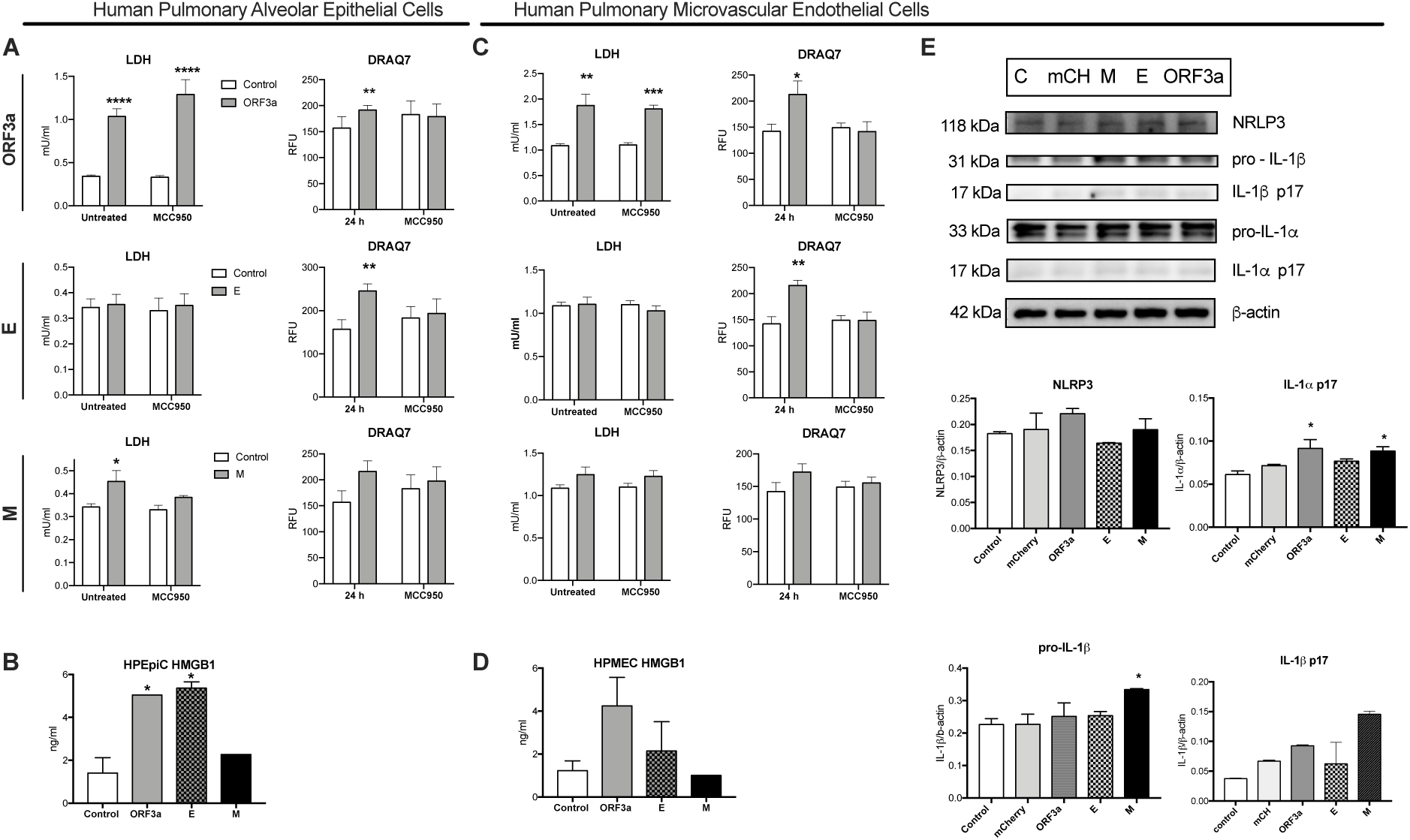
Effects of the ORF3a, E and M proteins expression the human primary pulmonary epithelial and endothelial cells. A. Assessment of cytotoxicity of the expressed viroporins in human pulmonary alveolar epithelial cells (HPAEpiCs) 24 hours after lentiviral infection by the LDH release (left column) and DRAQ7 incorporation (right column). B. Effects of the viroporins on HMGB-1 release by HPAEpiCs. C. Assessment of cytotoxicity of the expressed viroporins in human pulmonary microvascular endothelial cells (HPMECs) 24 hours after lentiviral infection by the LDH release (left column) and DRAQ7 incorporation (right column). D. Effects of the viroporins on HMGB-1 release by HPMECs. E. Immunoblots for the inflammasome proteins (upper panel) and their respective densitometric analysis (lower panel) analyzed in HPMECs. C- control, mCH-mCherry. Graphs represent means ± SEM, n=3-4. Two groups comparisons were analyzed with t-test. Multiple comparisons were performed with two-way ANOVA test. *p<0.05.

### SARS-CoV-2 activates inflammasome in human macrophages

The above described experiments revealed that specific viroporins are able to activate the inflammasome. To verify whether similar effects can be observed in more translational setting, we performed *in vitro* infection experiments of human monocyte-derived macrophages with SARS-CoV-2 to assess its potential to activate the inflammasome in controlled conditions. As shown in the **Figure 6 A**, human macrophages can be infected by SARS-CoV-2. Infection of human macrophages induced cleavage of IL-1 and GSDMD within eight hours post infection (**Fig. 6 B**). This was accompanied by the trending rise in LDH efflux and IL-18 that did not reach significance (**Fig. 6 C**). However, the infected macrophages did release both IL-1*β* and IL-33 (**Fig. 6 C**). These findings confirm that SARS-CoV-2 can activate inflammasome in infected macrophages.

**Figure 6.**
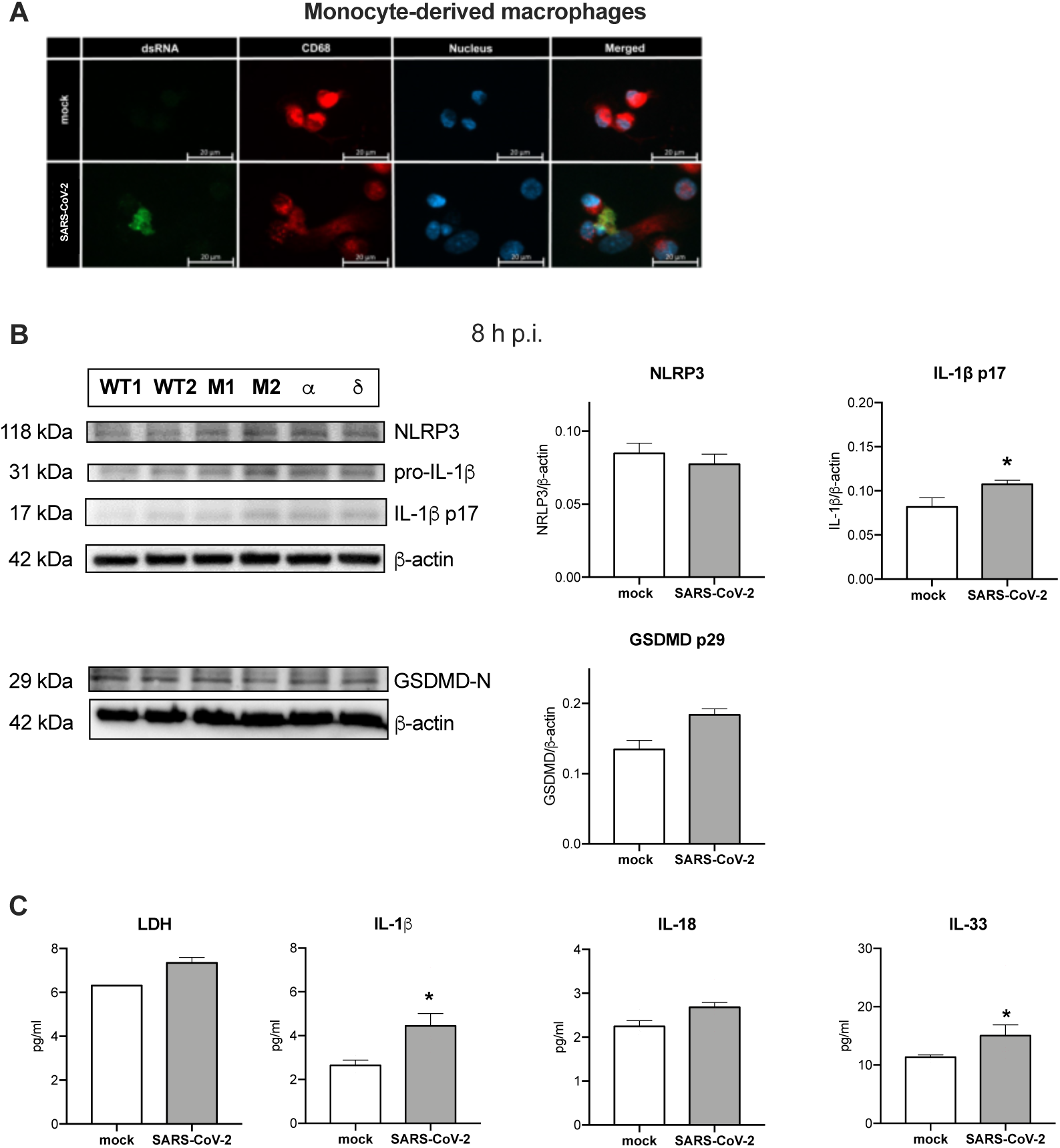
SARS-CoV-2 infection of monocyte-derived macrophages elicits inflammasome signaling. Human monocyte-derived macrophages were infected with SARS-CoV-2 at MOI 1 for 8 hours. A. Confocal microscopy analysis of the presence of SARS-CoV-2 dsRNA in CD68^+^macrophages. B. Western blot analysis of the expression of NLRP3, IL-1*β* (pro and cleaved p17), GSDMD-N and *β*-actin as loading control (left) upon infection with wild type, *α* and *δ* SARS-CoV-2 strains, M1, M2 – mock; densitometric analysis (right). C. Concentration of LDH, IL-1*β*, IL-18 and IL-33 was analyzed in the supernatant from the infected macrophages. Graphs represent means ± SEM, n=3-4. Two groups comparisons were analyzed with t-test. *p<0.05.

### Infection with SARS-CoV-2 induces hallmarks of inflammasome activation in alveolar epithelial and endothelial cells

Similarly, we tested the capacity of SARS-CoV-2 to activate inflammasome in the human primary pulmonary epithelial and endothelial cells. Firstly, we confirmed that in our setting SARS-CoV-2 infects the small airway epithelial cells (SAECs, **Fig. 7 A**). Then, we analyzed expression and cleavage of inflammasome-related proteins 1- and 3-days post-infection as the kinetic experiments showed slower dynamics of the activation of epithelial cells than macrophages. SAECs expressed the NLRP3 protein and at one day post-infection the level of cleaved IL-1*α* was increased (**Fig. 7 B**). Infected SAECs secreted IL-1*α*, IL-18 and IL-33 after one day and LDH release was observed after three days (**Fig. 7 C**). SARS-CoV-2 was also able to infect the pulmonary microvascular endothelial cells (HPMECs) (**Fig. 7 D**). NLRP3 protein was also present in HPMECs and infection with SARS-CoV-2 increased pro-IL-1*α* but had almost no effect on the GSDMD cleavage (**Fig. 7 E**). In accordance, the SARS-CoV-2 infection did not increase LDH release but led to the increased secretion of IL-1*α*, IL-18 and IL-33 (**Fig. 7 F**). Altogether, SARS-CoV-2 can induce inflammatory cell death in pulmonary epithelial but not endothelial cells and triggers the release of the inflammasome-related cytokines.

**Figure 7.**
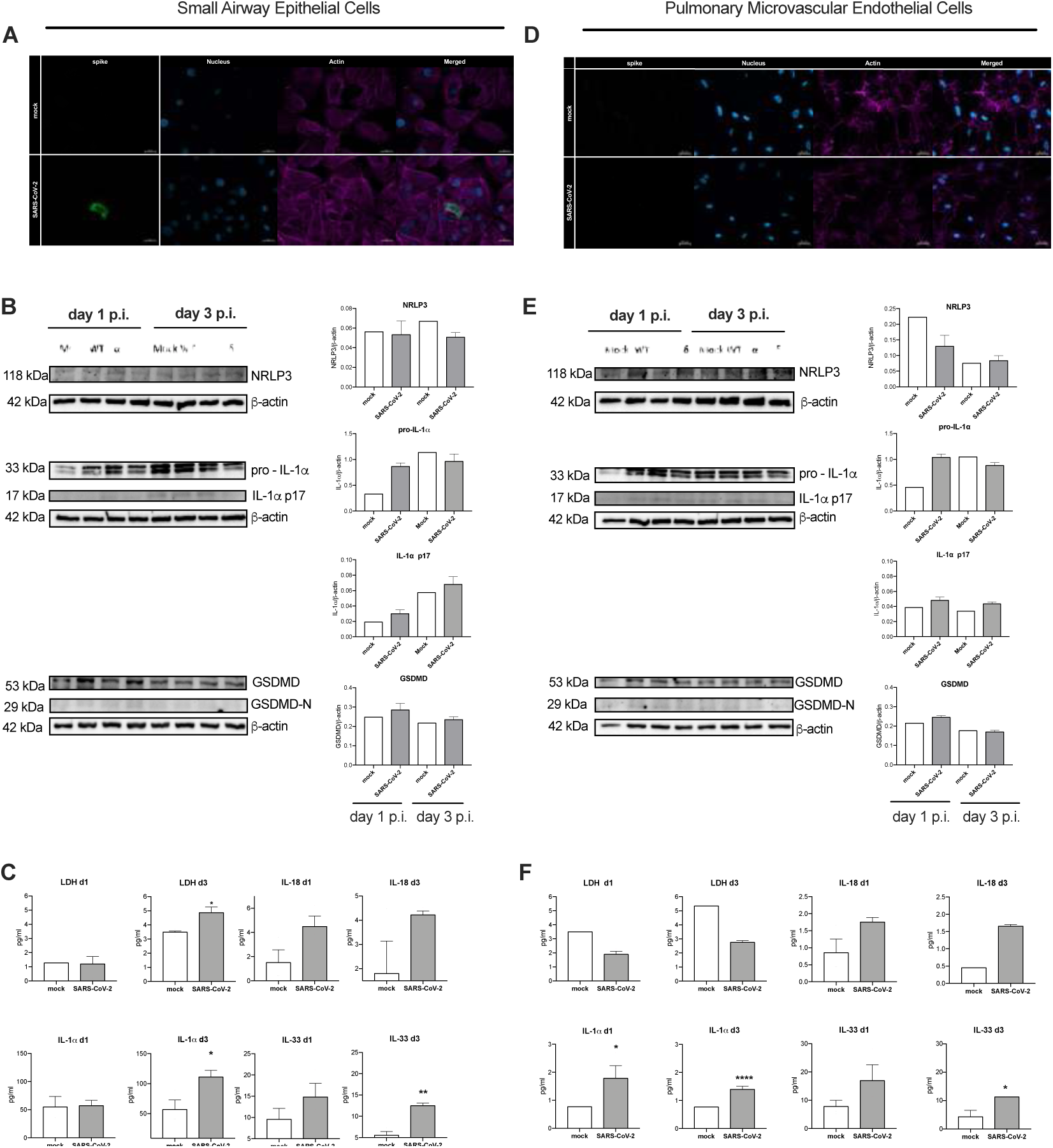
Effects of SARS-CoV-2 infection on human small airway epithelial cells (SAECs) and pulmonary microvascular endothelial cells (HPMEC) *in vitro*. A. Immunolocalization of spike protein in SAECs after 1 h of infection. B. Western blot analysis of the expression of NLRP3, IL-1*α* (pro and cleaved p17), GSDMD, GSDMD-N and *β*-actin as loading control (left) in SAECs upon infection with wilde type, *α* and *δ* SARS-CoV-2 strains; respective densitometric analysis (right column). C. Concentration of LDH, IL-18, IL- 1*α* and IL-33 was analyzed in the SAECs supernatant 1- and 3-day post infection. D. Immunolocalization of spike protein in HPMECs after 1 hour of infection. E. Western blot analysis of the expression of NLRP3, IL-1*α* (pro and cleaved p17), GSDMD, GSDMD-N and *β*-actin as loading control in HPMECs (left) upon infection with Wilde type, *α* and *δ* SARS-CoV-2 strains; respective densitometric analysis (right column). F. Concentration of LDH, IL- 18, IL-1*α* and IL-33 was analyzed in the HPMECs supernatant 1- and 3-day post infection. Graphs represent means ± SEM, n=3-4. Two groups comparisons were analyzed with t-test. *p<0.05.

## Discussion

SARS-CoV-2 infection triggers inflammasome signaling in the lungs of COVID-19 patients. This potent innate immunity mechanism was also suggested to be a driver of severe course of the disease [1, 23]. Although the potential of SARS-CoV-2 and its components to activate inflammasome was shown, a comprehensive study on the inflammasome signaling in the major sentinel cells of the alveolus, macrophages, epithelial and endothelial cells is missing. Here, we confirmed that SARS-CoV-2 viroporins ORF3a, E and M are capable of triggering NLRP3 inflammasome activation in human macrophages and inducing cell death in alveolar epithelial and endothelial cells. Moreover, we found that both viroporin stimulation and SARS-CoV-2 infection triggers IL-1*α* expression and cleavage in human pulmonary endothelial cells.

Despite conflicting reports on expression of ACE2 by human macrophages, we confirmed previous results showing the ability of SARS-CoV-2 to infect macrophages even in the absence of human IgGs [25]. Although the macrophage infection with SARS-CoV-2 was reported to be abortive [26] the viral molecules can induce cellular immunity. The use of time-lapse imaging of the reporter macrophage cell line enabled us to concomitantly trace the assembly of the inflammasome complex observed as ASC speck formation and lytic cell death. Notably, ORF3a and E proteins induced rapid inflammasome formation in THP-1 macrophages which was accompanied by IL-1*β* release. In addition, pyroptosis was also confirmed by the LDH release which was also found to be elevated in the blood of COVID-19 patients [27]. Using knock-out cell lines we showed that all investigated viroporins induced IL-1*β* maturation in NLRP3-dependent manner, while cell death was only partially reduced in the absence of this inflammasome. This suggests involvement of other cell death mechanisms such as caspase-8, AIM2 or PANoptosis [25, 28, 29]. Additionally, we evaluated the effects of the selected inhibitors on the viroporin-induced inflammasome activation. Several experimental readouts allowed us to observe differential effect of these inhibitors, e.g. MCC950, colchicine, disulfiram and KCl were able to reduce release of IL-1*β* induced by ORF3a, E and M (only two first listed inhibitors), while cell death was inhibited only by KCl (for ORF3a) and colchicine (for E protein). Intriguingly, only disulfiram reduced formation of ASC specks by protein E. The observed uncoupling of pyroptosis and IL-1*β* release inhibition can at least in part explain the lack of benefits with the use of colchicine or NLRP3 inhibitor in clinical trial [30, 31]. Zheng et al. [32] found that the extracellular protein E can prime the inflammasome (signal 1) via TLR2 but is insufficient to activate it (lack of signal 2). We found that when present intracellularly, the E protein can also activate inflammasome similarly to SARS-CoV protein E [14]. Noteworthy, in our experimental model we did not pre-stimulate macrophages with TLR agonists as lentiviral infection can induce NF-*κ*B signaling. Our results suggest that SARS-CoV-2 viroporins trigger uncoupled cell death and IL-1*β* release in human macrophages. As the lentiviral delivery of viral plasmids is a synthetic system, additionally we analyzed the responses of the monocyte-derived macrophages infected with SARS-CoV-2 virus and also observed IL-1*β* and gasdermin D cleavage. Infection with the complete virus complements the above-mentioned results as also structural proteins S and N and viral ssRNA were reported to be capable of inflammasome activation [16, 17, 33]. Furthermore, the mechanisms of inflammasome activation by intact virus involve TLR2 signaling and may be different from those elicited by viroporins [16].

As the airway epithelial cells are the primary target of SARS-CoV-2 infection, we first applied the real-time inflammasome activity monitoring with reporter bronchial endothelial cell line to trace responses induced by the expressed viroporins. Infection with lentivirus encoding all three tested proteins induced significant cell death in comparison to control lentivirus with ORF3a and E proteins causing the strongest effect which was partially reduced by NLRP3 inhibition with MCC950. In contrast, ASC speck assembly was not observed upon expression of neither of the viroporins. Likewise, primary human pulmonary alveolar epithelial cells which expressed viroporins showed hallmarks of pyroptosis and released HMGB1. Thus far, only ORF3a was reported to trigger pyroptosis in human alveolar epithelial cells [34]. Interestingly, neither of tested viroporins activated pro-apoptotic caspases-3/7 suggesting that other viral factors are involved in the activation of the NLRP1-caspase-3 pathway that can converge to PANoptosis in SARS-CoV-2 infected epithelial cells [18, 35]. Also, infection with SARS-CoV- 2 induced IL-1*α* expression and release, however this process was evident 3 days after infection. Infection with intact virus weakly triggered generation of the GSDMD-N form what stays in line with the recent finding that gasdermin E is the primary pyroptosis executor in lung epithelial cells [18]. Aside from pyroptosis, SARS-CoV-2 triggers RIP3-mediated necroptosis in epithelial cells which can also contribute to low level of the cleaved GSDMD-N [36]. The relatively low expression of cleaved IL-1*α* and low levels of secreted LDH could be explained by the rarity of the permissive AT-2 cells within the small airway epithelial cells. Simultaneously, SARS-CoV-2 induced release of IL-33, which belongs to IL-1 family and can be released via GSDMD pores [37]. High serum concentration of IL-33 is a marker of unfavorable COVID-19 outcome [38] and locally produced IL-33 was demonstrated in lungs with post-COVID-19 fibrosis patients [39]. Recently, Barnett, et al. [40] described a mechanism by which epithelial cells provide signal 2 to myeloid cells but do not activate inflammasome in response to SARS-CoV-2. In our hands infected epithelium also showed signs of membrane rupture and although did not release IL-1*β*, these cells released other IL-1 family cytokines. These findings support the major role of the inflammasome activation in epithelial cells in orchestrating the immune response during SARS-CoV-2 infection.

Some characteristic symptoms of COVID-19 are attributable to endotheliopathy [2] so we studied also the effects of SARS-CoV-2 viroporins on inflammasome activation in endothelial cells. Despite some reports showing infection of endothelial cells, we did not confirm the presence of SARS-CoV-2 which stays in line with others [41]. Yet, it can be speculated that variants like Omicron, which mostly utilizes TMPRSS2-independent endocytosis, could infect also endothelial cells [42]. Induced expression of ORF3a, E and M proteins triggered moderate cell death in immortalized HUVEC cell line with no signs of ASC specks assembly. However, MCC950 reduced cell death in the ORF3a and M protein treated cells. The primary microvascular pulmonary endothelial cells showed hallmarks of lytic cell death upon viroporin expression. Interestingly, ORF3a also activated apoptotic caspases. ORF3a and M proteins induced cleavage of IL-1*α* and the latter also IL-1*β*. Treatment with SARS-CoV-2 intact virus evoked also IL-1*α* expression and release together with IL-33. The role of endothelial cells in IL-1*α* production during SARS-CoV-2 has not been reported to our knowledge and it has important consequences. IL-1*α* was shown to promote pathological myeloid cell expansion in the pulmonary infections [43] and cross-activate the coagulation system [44, 45]. This could add another player into the immunopathogenesis of pulmonary microthrombosis, which is a unique feature of severe COVID-19 [2]. Together, with epithelial cells, IL-1*α* expressing endothelium could explain positive outcome of anakinra (IL-1 receptor antagonist) in a subgroup of COVID-19 patients [10], and specifically IL-1*α* blockade in animal model of COVID-19 [46].

There are several limitations of our studies. Firstly, we used lentiviral expression system to deliver specific viroporins to target cells which reflects actively replicating rather than abortive viral infection. Yet, such an approach allows to dissect responses to single viral proteins. Although we performed experiments with SARS-CoV-2 infections we were not able to manipulate genes encoding viroporins.

In summary, we revealed that ORF3a but also E and M SARS-CoV-2 proteins are capable of activating the inflammasome in human macrophages and pulmonary epithelial and endothelial cells. Heterogenous effects of the inflammasome inhibitors on pyroptosis and IL-1 release can explain the poor clinical efficacy of these drugs in clinical trials. Importantly, both single viroporins and SARS-CoV-2 virus could induce IL-1*α* and IL-33 expression and release not only in pulmonary epithelial cell but also microvascular endothelium shedding novel light on the inflammatory response of endothelium in COVID-19.

## Supporting information

Supplementary results

## Acknowledgements

not applicable.

## Funding

This work was funded by the Polish National Science Centre grant no: UMO- 2020/01/0/NZ6/00218 (COVID-19) and the BMBF, funding program Photonics Research Germany (13N15745) and is associated into the Leibniz Center for Photonics in Infection Research (LPI) to SDE.

## Conflict of interest

The authors declare that they have no competing financial interest or personal relationships that could appear to influence the work reported in this paper.

## Authors

Conceived and designed experiments: TS, EK, SDE. Performed the experiments: MAL, PK, GH, MB, KH, DN, MG, LS. Analyzed the data: MAL, SDE, TS. Wrote the paper: MAL, PK, KG, APW, SDE, EK, TS. Acquire funding: SDE and TS.

## Data availability statement

Original data are available upon request.

## Notes

### Competing Interest Statement

The authors have declared no competing interest.

