## Supplementary results for "SARS-CoV-2 and its ORF3a, E and M viroporins activate inflammasome in human macrophages and induce of IL-1*α* in pulmonary epithelial and endothelial cells"

**Supplementary Figure 1. Validation of the novel inflammasome reporter cell lines.** Immortalized cell lines expressing the ASC::GFP protein were stimulated with LPS and nigericin. A. human bronchial epithelial cells -3 KT cell line. B. human umbilical vein endothelial cells/TERT cell line.

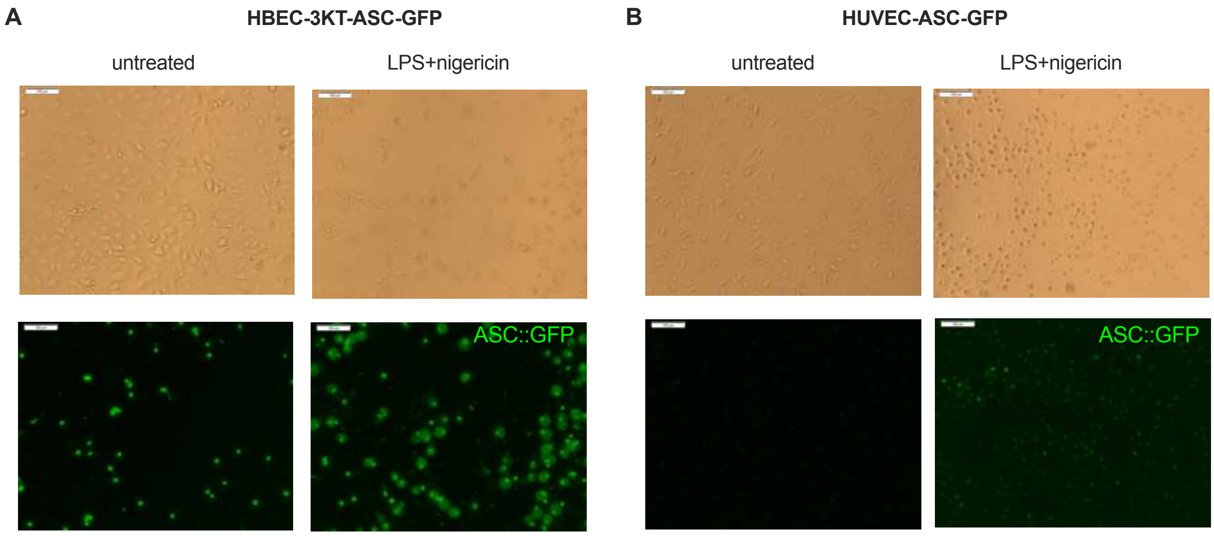


**Supplementary Figure 2. Analysis of the activation of apoptotic caspases in human pulmonary epithelial cells and human pulmonary microvascular endothelium by the viroporins** A. Activation of caspase-3/7 in HPAEpiCs by the viroporins; representative pictures (left) and fluorometric readouts (right) are shown. F. Activation of caspase-3/7 in HPMECs by the viroporins; representative pictures (left) and fluorometric readouts (right) are shown. Graphs represent means ± SEM, n=3-4. Two groups comparisons were analyzed with t-test. Multiple comparisons were performed with two-way ANOVA test. *p<0.05.

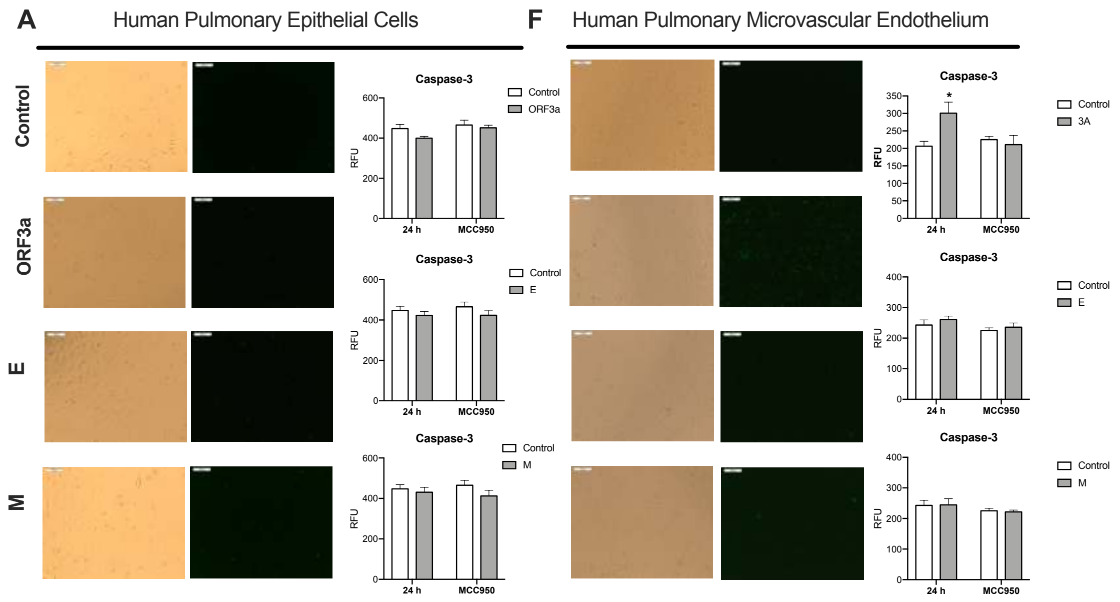
